# Exploring natural allies: Survey and identification of larval parasitoids for sustainable grape berry moth management in vineyards

**DOI:** 10.1101/2024.12.27.630474

**Authors:** Jesus H. Gomez-Llano, Neetu Khanal, Flor E. Acevedo

**Author notes:** These authors contributed equally to this work.

## Abstract

The American grape berry moth (GBM), *Paralobesia viteana* Clemens (Lepidoptera: Tortricidae) is an economically important pest of grapes. The larvae of this insect burrow inside the fruit upon hatching, consuming, and contaminating grapes and clusters. Current GBM management relies on pesticide applications, which do not offer complete protection due to the cryptic behavior of the larvae and asynchrony in egg-laying, highlighting the need to develop new management strategies. In this study, we identified GBM larval parasitoids in commercial vineyards and quantified their parasitism rates. Parasitoid samplings were conducted biweekly in six conventionally managed Concord vineyards in Erie County, Pennsylvania, during the 2023 and 2024 growing seasons. GBM-infested samples were monitored daily to track the emergence of both parasitoids and GBM, enabling the calculation of parasitism rates. We identified eight parasitoid species: *Enytus obliteratus*, *Campoplex tortricidae*, *Scambus* sp, *Glypta depressa* cf, *Glypta ohioensis* cf, and *Glypta ignota* cf. (Hymenoptera: Ichneumonidae); *Bracon variabilis* (Hymenoptera: Braconidae), and *Goniozus fratellus* (Hymenoptera: Bethylidae) praying on GBM larvae. From these, *B. variabilis*, *E. obliteratus*, and *G. fratellus* were the most abundant. We also designed a graphic taxonomic key to facilitate the identification of these species. The parasitoid abundance differed over the growing season but was greatest in early August, reaching parasitism rates of up to 39% and 52.1% in 2023 and 2024, respectively. Our results demonstrate that GBM has several larval parasitoids that help reduce its populations in commercial vineyards. This project represented a first step toward our understanding of the GBM native natural enemies present in the Lake Erie Region and their potential use in management programs.

## Introduction

The grape berry moth (GBM), *Paralobesia viteana* is a highly destructive pest of grapes in eastern North America [1]. The larvae of this insect feed on grape clusters causing mechanical injury and increasing the vulnerability to pathogen infection ultimately, resulting in yield losses and a decrease in juice and wine quality. Management of GBM currently relies on insecticide sprays timed using a degree-day model developed by Tobin et al. [2]. Although insecticides confer some protection against this insect, their constant use poses human and environmental risks and can lead to the development of GBM resistance to these products [3]. With the long-term goal of developing more sustainable strategies for GBM control, this study focused on the identification of larval parasitoids in Concord (*Vitis labrusca*) vineyards in northwestern Pennsylvania (U.S.A).

GBM is native to eastern North America and has coevolved with numerous natural enemies in its original habitat. Studies in the Finger Lakes (New York), the Lake Erie Region (New York and Pennsylvania), and Michigan have identified several egg and larval parasitoids of GBM (Table 1). However, these studies date back several decades underlining the need for new surveys to identify current GBM natural enemies that have endured increases in farming land, recurrent pesticide sprays, and climate changes in grape-growing regions. In a 2-year survey conducted in the Finger Lakes in 1986-87, the natural community of egg and larval GBM parasitoids caused 12-42% mortality throughout the growing season [4]. The most abundant parasitoid in this study was *Trichogramma pretiosum* Riley to which a large percentage of GBM mortality (20.2%) was attributed. Studies conducted in Michigan vineyards reported *Sinophorus* sp. as the most abundant parasitoid causing 11-76% GBM mortality in field conditions [5]. Surveys in Arkansas vineyards found GBM parasitism of 3.6 - 48.6% from Ichneumonid and Braconid parasitoids with no detail of the species reported [6]. Although some work has been conducted in the identification of GBM natural enemies, their efficacy in reducing insect infestations in field conditions remains untested except for one egg parasitoid species, *Trichogramma minutum* Riley. *T. minutum*, is a native parasitoid that was found infesting GBM eggs in northwestern Pennsylvania, but its natural parasitism rates were very low to decrease pest populations [7]. Inundate releases of *T. minutum* in commercial vineyards showed a significant decrease in GBM damage [8], demonstrating the potential of natural enemies for controlling this pest.

**Table 1.**
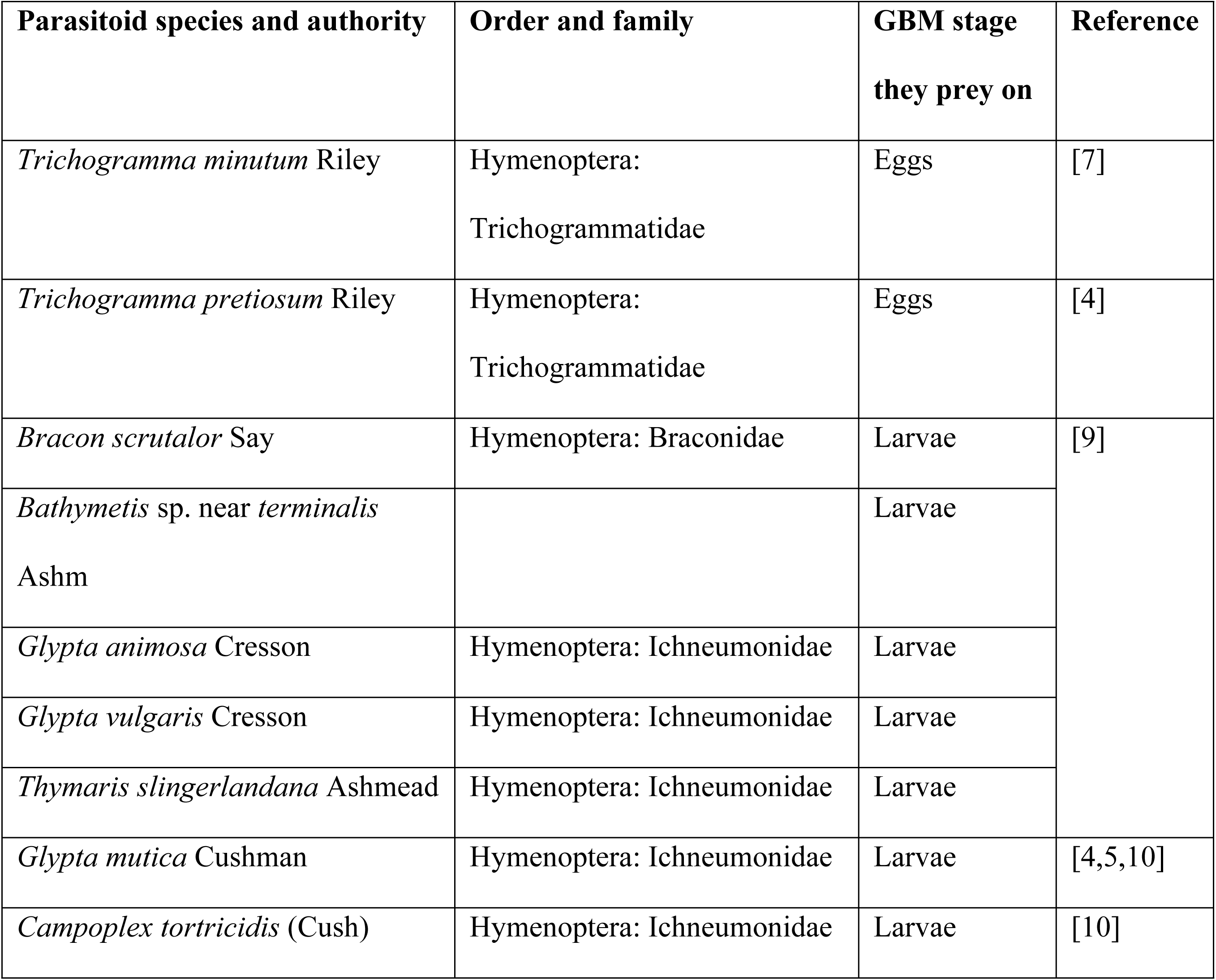
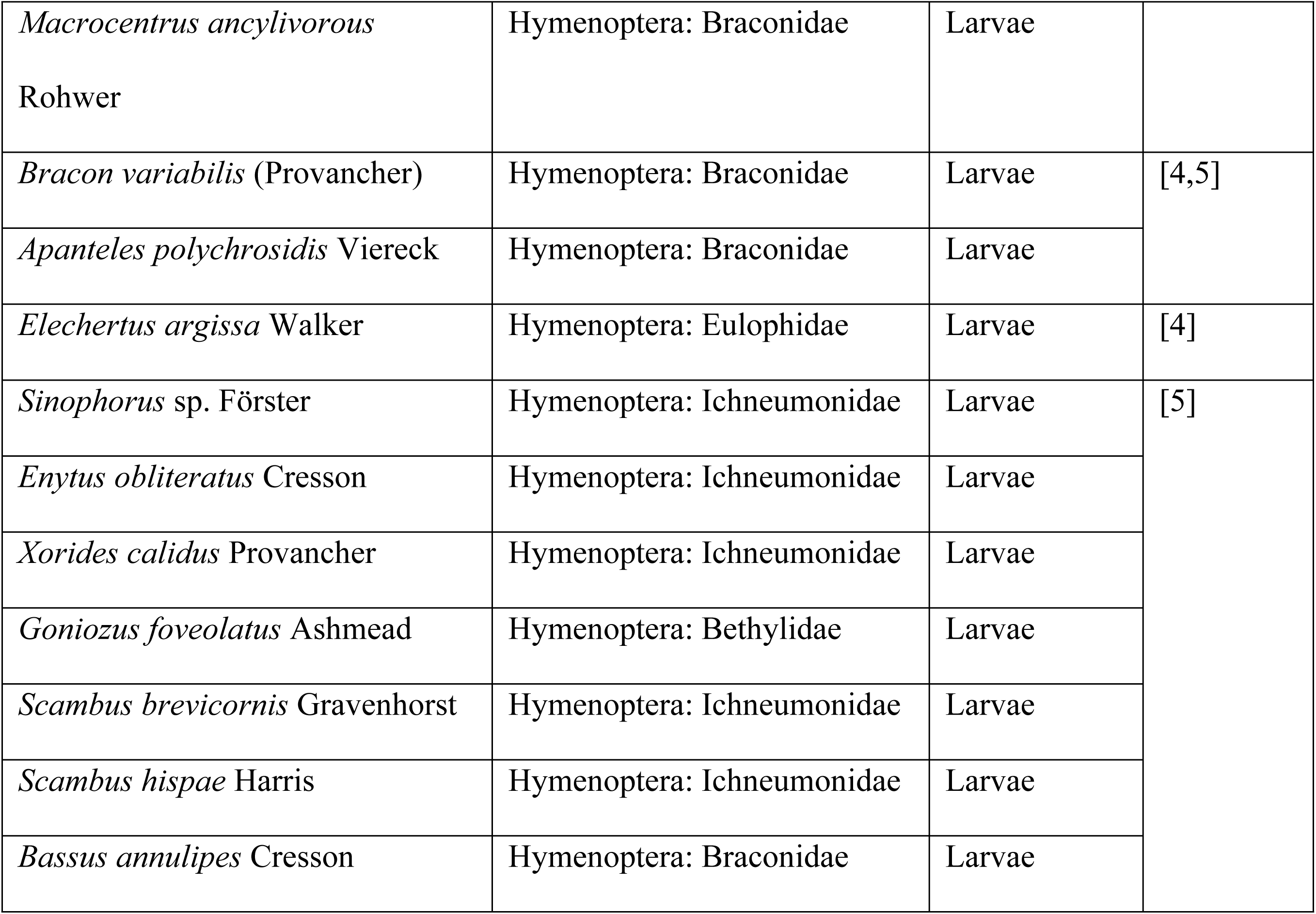
Grape berry moth parasitoids previously identified in U.S. vineyards.

The biology and ecology of GBM, make the control of this insect particularly challenging. GBM evolved with wild grape hosts and became a pest in vineyards established east of the Mississippi River [1]. Adults typically emerge from overwintered pupae between May and June [11,12].

However, the termination of diapause is variable lasting about six weeks [12], resulting in asynchronous egg laying over the growing season. Mated females oviposit on developing buds, flower clusters, or berries from wild and cultivated grapevines [1,13]. Females lay an average of 60 eggs in their lifetime from which more than 86% successfully hatch in controlled conditions [14]. Larvae emerge after 3-4 days and create webs in the cluster [1,11]. Early in the season, larvae feed on buds or flower clusters, whereas later in the season, they burrow into the fruit within hours of egg-hatch. Inside the grape berries, larvae remain protected against chemical control while consuming the internal tissue of the berry. The larval period takes an average of 18.5 days at 25 ± 1 °C [14]. After fully developed, fourth instar larvae exit the berry and pupate by cutting a flap in a grapevine leaf and forming a silken chamber [11,12]. The pupa stage takes an average of 9.52 days to develop at 25 ± 1 °C, and the longevity of adults is on average 15.6 days in controlled conditions [14]. The oviposition period is 7.3 days on average, with some variation associated with the grape cultivar [14]. The number of GBM generations per season varies with temperature accumulation; there are two to three generations in the Lake Erie region and central New York State [1,15–17], whereas, in southern Missouri and Arkansas, there can be up to four generations [1,18]. In the Lake Erie region, the first GBM generation develops in wild grapes (*Vitis* spp), and subsequent generations develop in cultivated grapes, where they cause significant crop losses [7]. Access to widespread wild hosts, the cryptic feeding habit of the larvae, the prolonged oviposition period of females, and the large variation of diapause termination contribute to the challenges of controlling this insect.

The variation in diapause termination results in constant GMB pressure that reduces the effectiveness of chemical control that targets peaks of egg laying [12]. Therefore, a long-lasting control strategy that reduces pest populations over several weeks would be ideal. Biological control using egg and larval parasitoids holds great potential for controlling GBM because they may have the capability of controlling the pest for several days. Additionally, parasitoids may be able to control GBM in both wild and cultivated grapes. Augmentative releases of parasitoids have great potential to reduce GBM populations below economic injury levels [7]. Furthermore, parasitoids could offer long-lasting pest control if they successfully establish in the field, but research is needed to determine the feasibility of this strategy.

In this study, we surveyed and assessed the parasitism rates of GBM larval parasitoids in six vineyards in Erie County (Pennsylvania) throughout the 2023 and 2024 growing seasons. We also developed a graphical taxonomic key to facilitate further identification of these species. Our results provide knowledge of the current diversity of GBM larval parasitoids in commercial vineyards of northwest Pennsylvania and provide the foundation for future studies to test the effectiveness of the identified species on GBM control in field conditions.

## Materials and methods

### 1. Sampling sites and specimen collection from the field

Surveys were conducted from June to September 2023 and 2024 in six conventional vineyards located in Erie County, northwest Pennsylvania. Among these six sites, four (Site 1: 42°11’20.0"N, 79°53’05.6"W; Site 2: 42°12’29.0"N, 79°53’26.6"W; Site 3: 42°12’30.9"N, 79°53’31.7"W, and Site 4: 42°13’30.1"N, 79°49’10.5"W) were situated in North East, Pennsylvania, to the east of Erie city, while the remaining two sites (Site 5: 42°02’04.7"N, 80°18’20.9"W, and Site 6: 42°01’55.0"N, 80°17’20.0"W) were located to the west of Erie city (Fig 1). These vineyards were selected for their recurrent history of GBM infestation and were all planted with *V. labrusca* cv. Concord. These vineyards also had at least one border near a deciduous wood area and received insecticide applications according to each grower’s standard insect control program. Grapevine blossoms and grapes with signs of GBM infestation (blossoms or grapes attached with silk and purple-colored grapes with holes) were collected from the six sites throughout the growing season in both years. All samples were collected from vineyard borders as previous research indicates that borders are more heavily infested with GBM and, consequently, have a greater number of natural enemies [4,5]. The first sampling in 2023 (June 15) was from wild grapevines growing at the borders of the cultivated plots and the subsequent samples were obtained from the adjacent commercial Concord vineyards. During the first sampling, we collected several bunches of wild grape blossoms, while in the subsequent samplings, we collected ∼ 100 - 150 grapes per site from the vineyard borders adjacent to wooded areas. In 2024, we attempted to collect samples from wild grapes in early June, but there were too few insects from only a few sites. Therefore, all our samplings for this year were from commercial vineyards.

**Fig 1.**
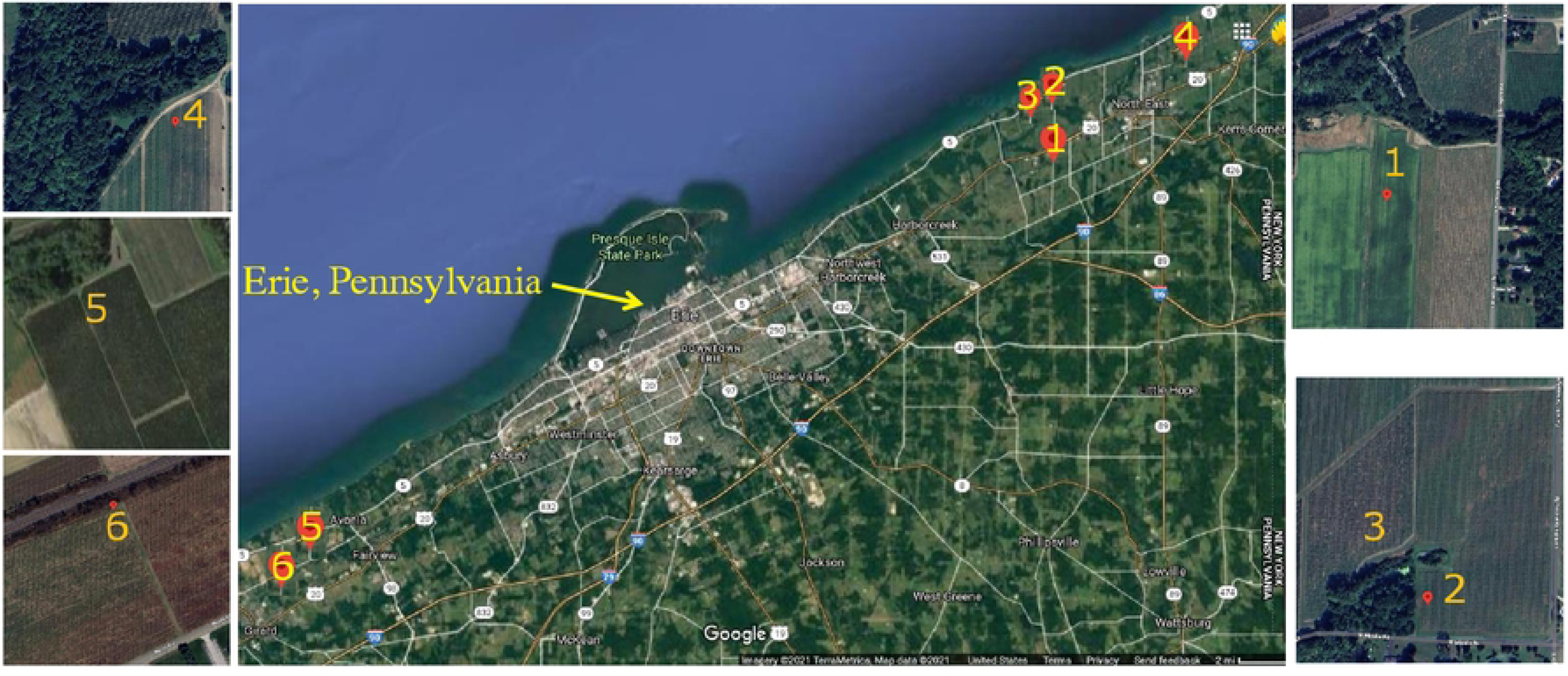
Sampling sites of grape berry moth larval parasitoids in Northwest Pennsylvania. A closeup of each vineyard is presented outside the map with a matching number of each sampling site 1-6.

### 2. Specimen rearing in laboratory conditions and calculation of parasitism rates

Field-collected specimens were taken to a laboratory located in Penn State Behrend (Erie, PA) for rearing and collection of GBM larval parasitoids. The samples were kept in a Conviron growth chamber (GEN200SH) at a temperature of 25 ± 1 °C, 70% ± 2% relative humidity (RH), and a photoperiod of 16:8 h (L:D). Grape blossoms from wild grapes were inspected under a stereoscope (Leica S9D Nuhsbaum Inc, McHenry Il, USA) to carefully remove GBM larva using a soft fine paintbrush; each larva was individualized in 1 oz plastic cups (Dart Conex Complements 301100PC, WebstaurantStore, Landcaster PA) and fed with wild grapes collected from the field. Each cup was provided with a piece of paper towel (∼2 squared cm) as a substrate for GBM pupation. The cups were inspected periodically to record moth or parasitoid emergence. GBM-infested grapes were also taken to the lab and placed in 450 ml plastic containers (6 cm tall, 11 cm in diameter) covered at the top with a fine fabric mesh fixed with a rubber band. Pieces of paper towel (∼2 squared cm) were placed inside the containers as substrate for GBM pupation. The cups were periodically inspected for GBM or parasitoid pupae as well as for adult emergence. The number of GBM and the number of parasitoids was recorded for each site and sampling time.

### 3. Parasitoid identification and development of a taxonomic key

Adult parasitoid specimens were stored in 1.5 ml plastic tubes either dry at -20°C or in 70% ethanol. These specimens were identified at the genus and species level using taxonomical keys for the families Ichneumonidae[19–22], Bethylidae [23], and Braconidae [24,25]. In addition, entomological material from the Frost Entomological Museum at Penn State University (State College, PA), the Cornell University Insect Collection (Ithaca, NY), and the USDA Systematic Entomology Laboratory (Smithsonian Institution, Washington D.C.) were reviewed to confirm the taxonomic identification of the specimens. Voucher specimens are stored in the Frost Entomological Museum, and extra samples are kept at Flor Acevedo’s laboratory. Lastly, we developed a taxonomic key following the terminology proposed by Townes [19], Evans [23], and Achterberg [24], to facilitate the identification of the primary parasitoids documented in this research. Images of parasitoid specimens were taken under a Leica S9D stereomicroscope with an attached digital camera Canon EOS 90D (Melville, NY). Images of small specimens and body structures were taken with an FEI quanta 650 FEG Environmental Scanning Electron microscope (SEM).

### Data analyses

We calculated the percent of parasitism in the samples collected from the field for each sampling site and sampling date in 2023 and 2024. The parasitism rates were expressed as the number of parasitoids divided by the sum of parasitoids and GBM stages that emerged multiplied by 100.

The effects of the different sites and sampling times on the mean percent of parasitism were assessed separately for each year using a beta regression model with a logit link function for the mean (*mu*) and an identity link function for the dispersion parameter (*phi*) using the response variable as a proportion between 0 and 1. We used *phi* coefficients to determine the variability of the data and log-likelihood to determine how well the model fitted the data [26]. With these methods, we tested the null hypothesis (H_0_) of no differences in the percent of parasitism between sites and sampling times.

The parasitoid community composition in the sampling sites for each year was assessed by calculating the diversity indexes of Shannon-Weaver and Simpson using the “vegan” R package [27]. Statistical differences in the Shannon-Weaver diversity indexes between sites were assessed with Hutcheson’s t-test [28]. All statistical analyses and graphs were made using R [29].

## Results

### Field parasitism rates

We collected a total of 113, and 159 GBM larval parasitoid specimens from all sites and samplings carried out in 2023 and 2024, respectively. In 2023, the highest parasitism rates were observed in the first sampling of the season in wild grapes (30.95% on average; *Z* = -5.4, *P* < 0.001), however, the number of samples with GBM living stages was very few in this sampling. The second highest parasitism rates were found in the sampling of Aug 7 with an average of 16.4% across sites while the lowest was found on July 11 with 2.7%. Site 1 had the highest parasitism across sampling dates with an average of 21.7% (*Z* = -3.4, *P* < 0.001), followed by site 6 with 13.6% and site 2 with 12%. Site 3 had the lowest parasitism rates with an average of 3.4% (Fig 2A; S1 Table). In 2024, the percentage of parasitism between sampling times was not statistically different (*P* > 0.05). Descriptively, the average highest parasitism rate across sites was 16.4% in the sampling conducted on Aug 5^th^, followed by the sampling conducted on July 23 (15.27%). The average lowest parasitism rates were found on Jun 24, with 1.6%. Site 1 had the highest parasitism across sampling dates, with an average of 19.3% (*Z* = -4.2, *P* < 0.001 followed by site 4, with 9%, and site 2, with 8%. Site 5 had the lowest parasitism rate with 3% (*Z* = -2.37, *P* < 0.05; Fig 2B; S2 Table).

**Fig 2.**
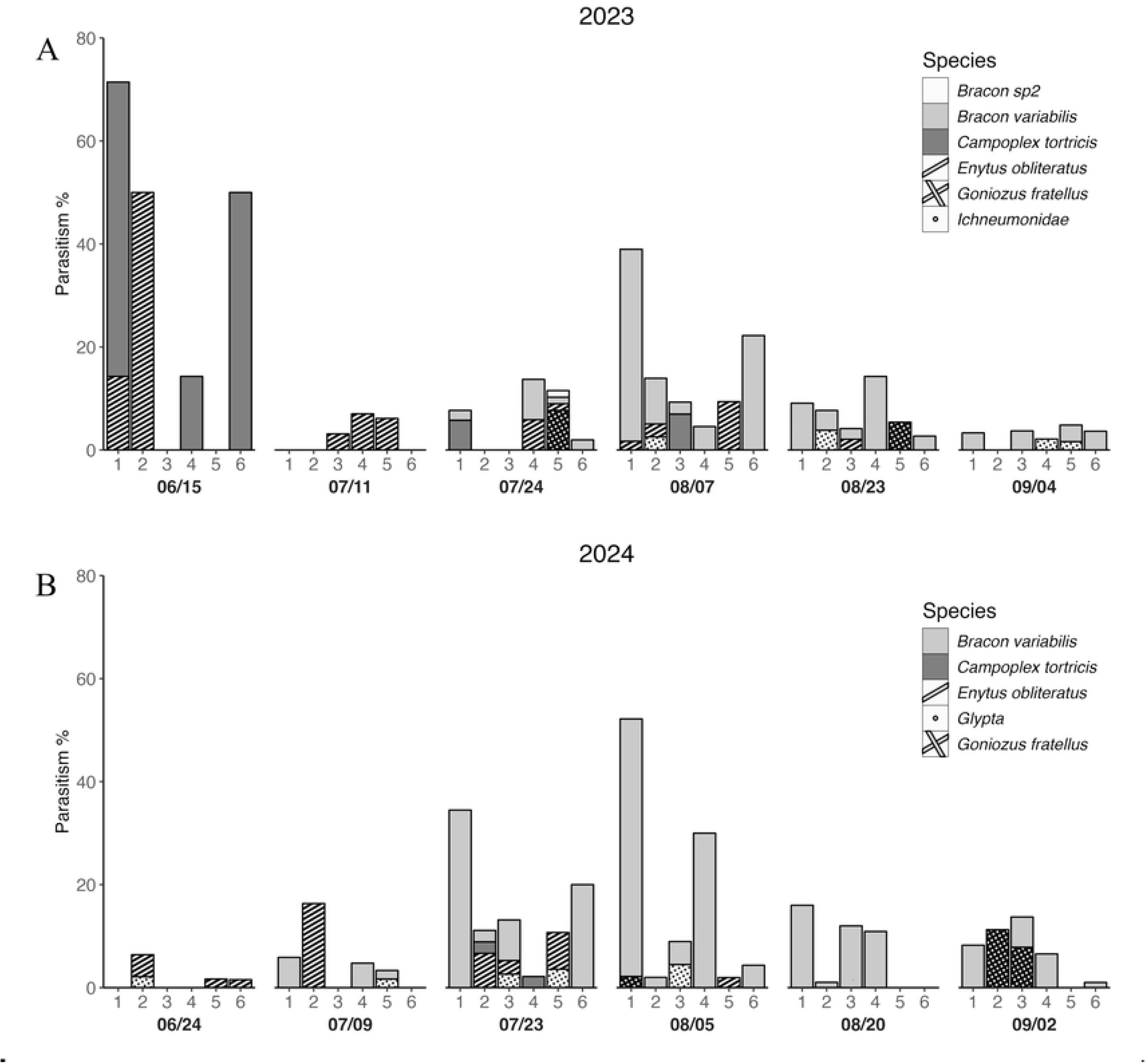
Field parasitism rates of GBM larval parasitoids. Percent of parasitism of GBM larval parasitoids in six different sites (1-6) and sampling times over the growing seasons of **(A)** 2023 and **(B)** 2024.

### Parasitoid identification and abundance

In our survey, we identified a total of 8 parasitoid species belonging to three Hymenoptera families. The identified species were *Enytus obliteratus*, *Campoplex tortricidis, Scambus* sp, three species of the genus *Glypta* (Hymenoptera: Ichneumonidae): *Glypta depressa* cf, *Glypta ohioensis* cf, and *Glypta ignota* cf., *Bracon variabilis* (Hymenoptera: Braconidae), and *Goniozus fratellus* Evans (Hymenoptera: Bethylidae). Additionally, we found three morphospecies within the *Scambus* genus, and one within *Glypta* that could not be identified further due to the lack of taxonomic keys. From these, *B. variabilis* and *G. fratellus* are ectoparasitoids, and the others are endoparasitoids. We collected the same parasitoid species in 2023 and 2024, except for *Scambus* sp., which was absent in our 2024 samplings. The most abundant parasitoid species are depicted in Fig 3.

**Figure 3.**
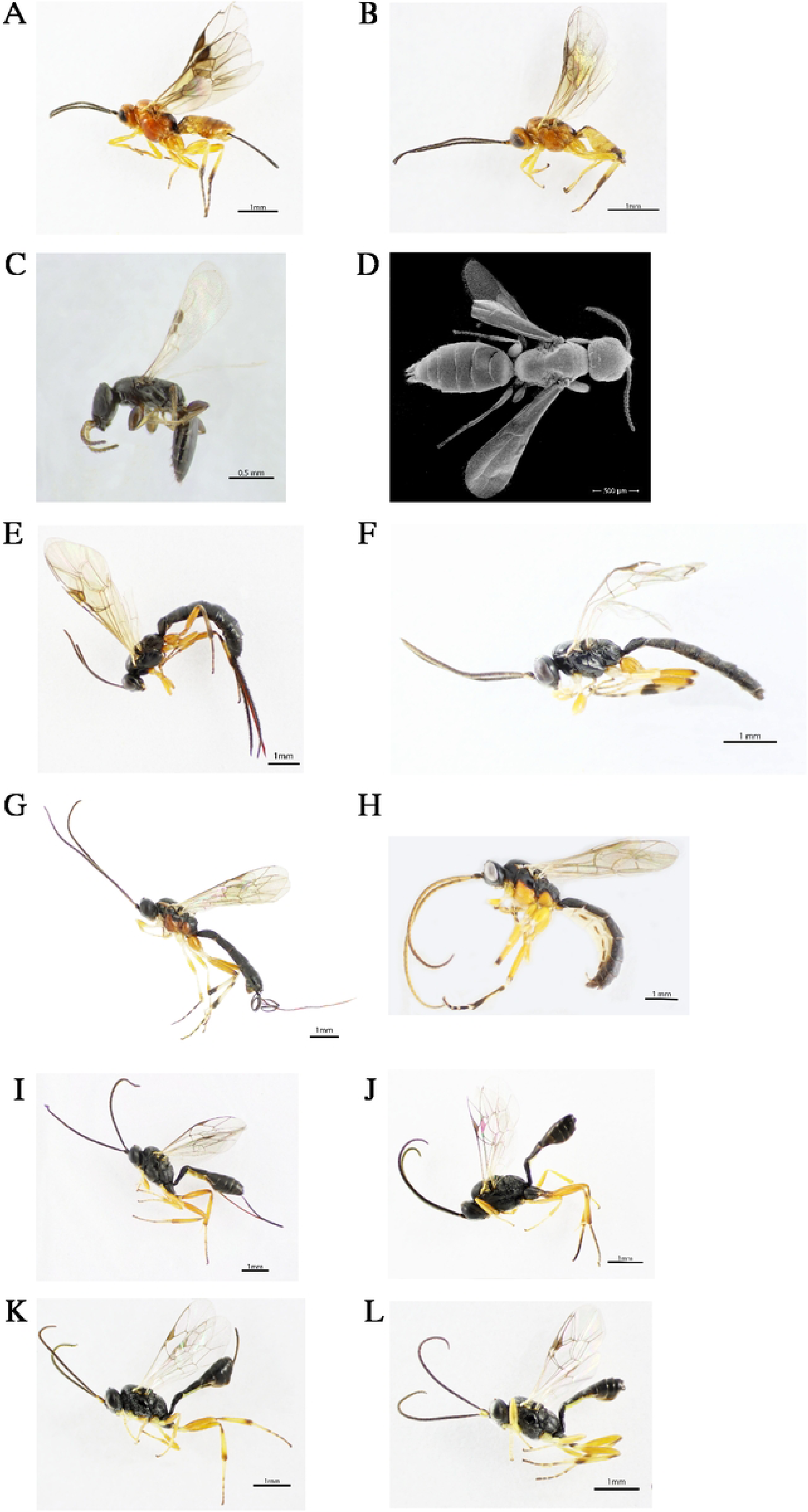
Grape berry moth larval parasitoids identified in northwest Pennsylvania. (**A-B**) *Bracon variabilis*, **(C-D)** *Goniozus fratellus,* **(E-F)** *Scambus* sp., **(G-H)** *Glypta* sp., **(I-J)** *Campoplex tortricidis*, **(K-L)** *Enytus obliteratus*.

The abundance of these parasitoids in the field varied throughout the growing season. In 2023, the most abundant parasitoid species were *B. variabilis* and *E. obliteratus*, comprising 56.6% and 18.6% of all parasitoids found, respectively (Fig 2A). In 2024, *B. variabilis* was also the most abundant species, representing 68.6 % followed by *G. fratellus* and *E. obliteratus* comprising 13.8% and 11.9%, respectively, of all parasitoids found (Fig 2B). *B. variabilis* was recovered from samples collected from July to September, whereas *E. obliteratus* was only present from June to mid-August. Notably, *G. fratellus* was only found late in the season (from late July to September). The remaining species were randomly found in samples collected throughout the season (Fig 2).

There were also differences in the parasitoid community composition in the sampling sites. In 2023, site 5 had the highest diversity compared with sites 1 and 6 (*P* < 0.001), but was no different from sites 2, 3, and 4 (*P* > 0.05) (Table 2, Fig 4A). In 2024, sites 2, 3, and 5 had the highest diversity compared with sites 1 and 4 (*P* < 0.001). Site 2 was also different from site 6 (*P* < 0.005) (Table 2, Fig 4B). In general, sites 2 and 5 were the most diverse during the two years of the study (Table 2, Fig 4).

**Fig 4.**
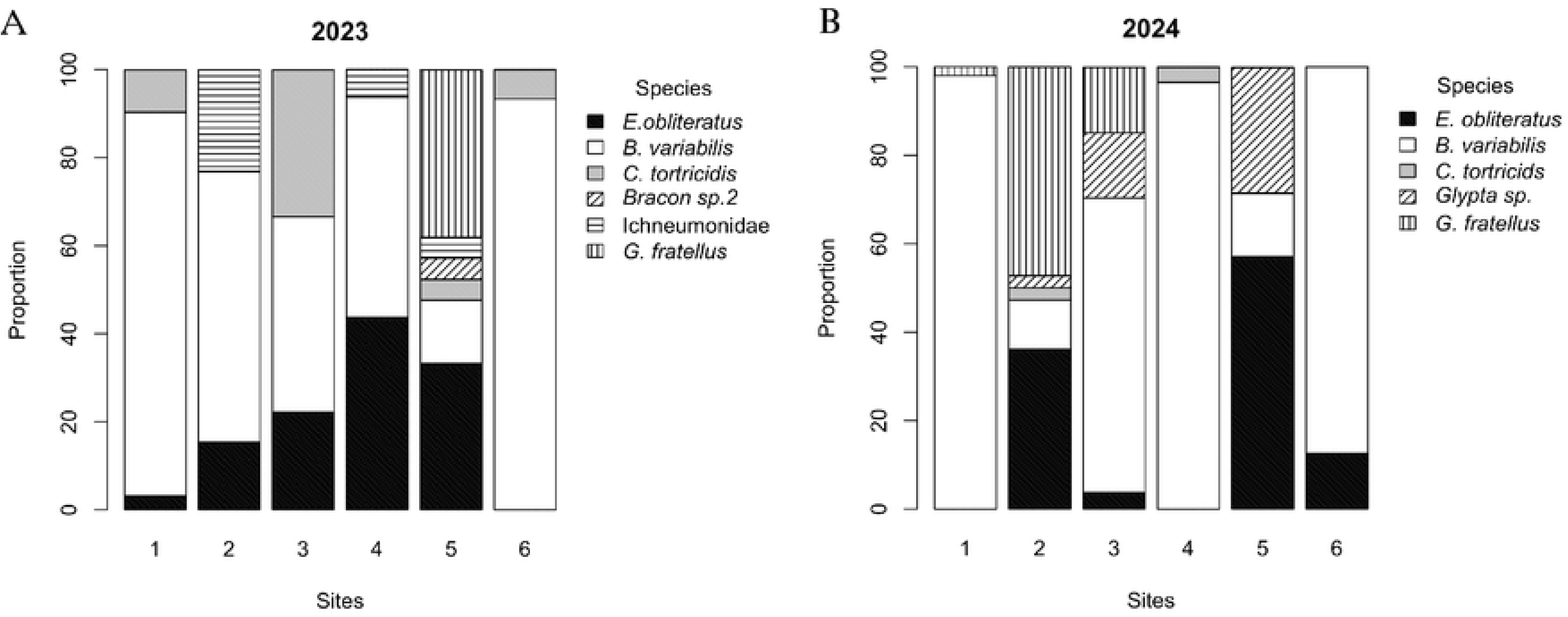
Abundance of GBM larval parasitoids across six different sites in (A) 2023 and (B) 2024.

**Table 2.**
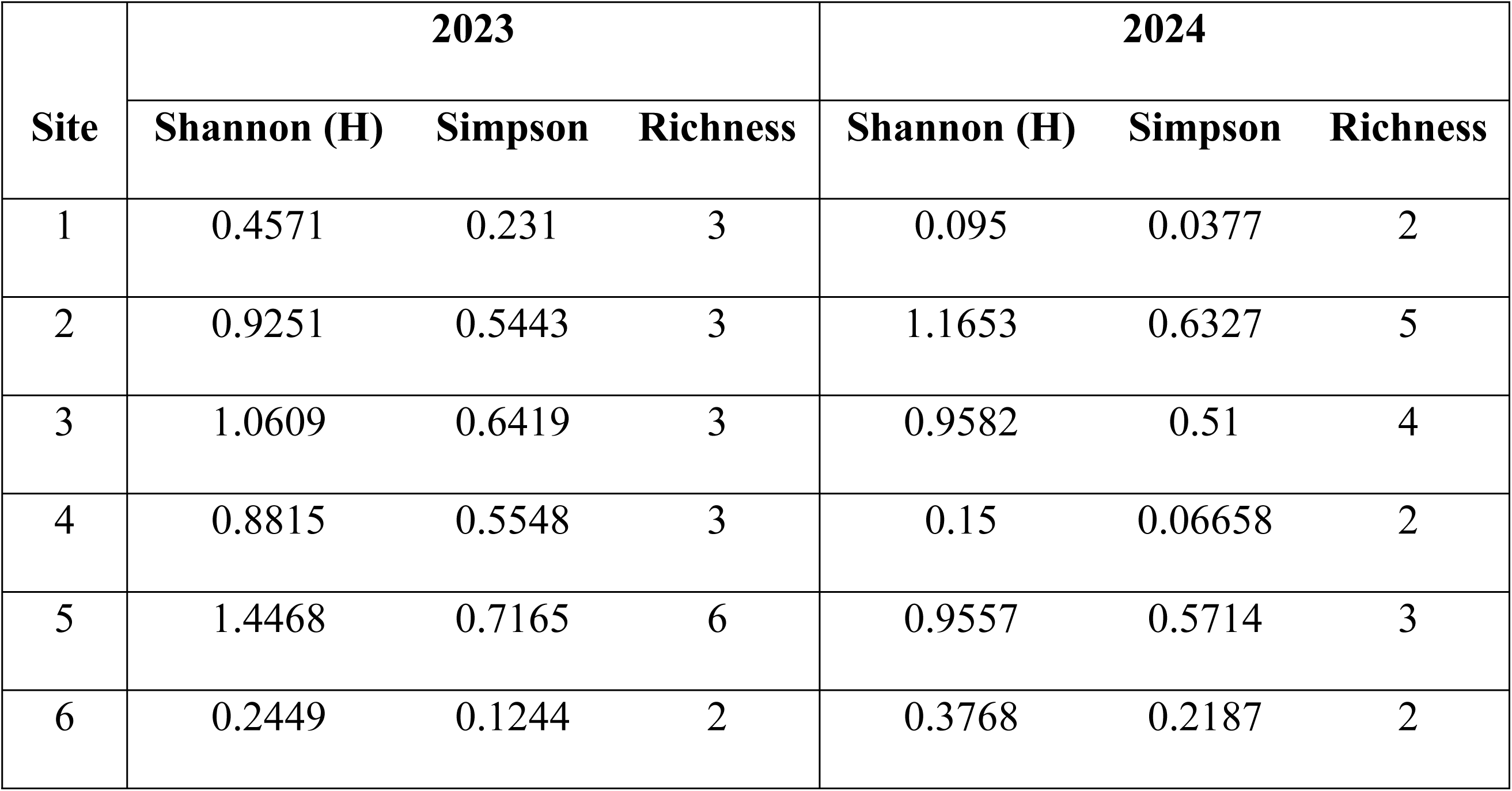
Alpha diversity indexes for each vineyard site (1-6) during 2023 and 2024.

### Taxonomic key for the GBM parasitoids identified in this study

1a. Fore wing vein 2m-cu present (Figs 5A-B) 3

**Figure 5.**
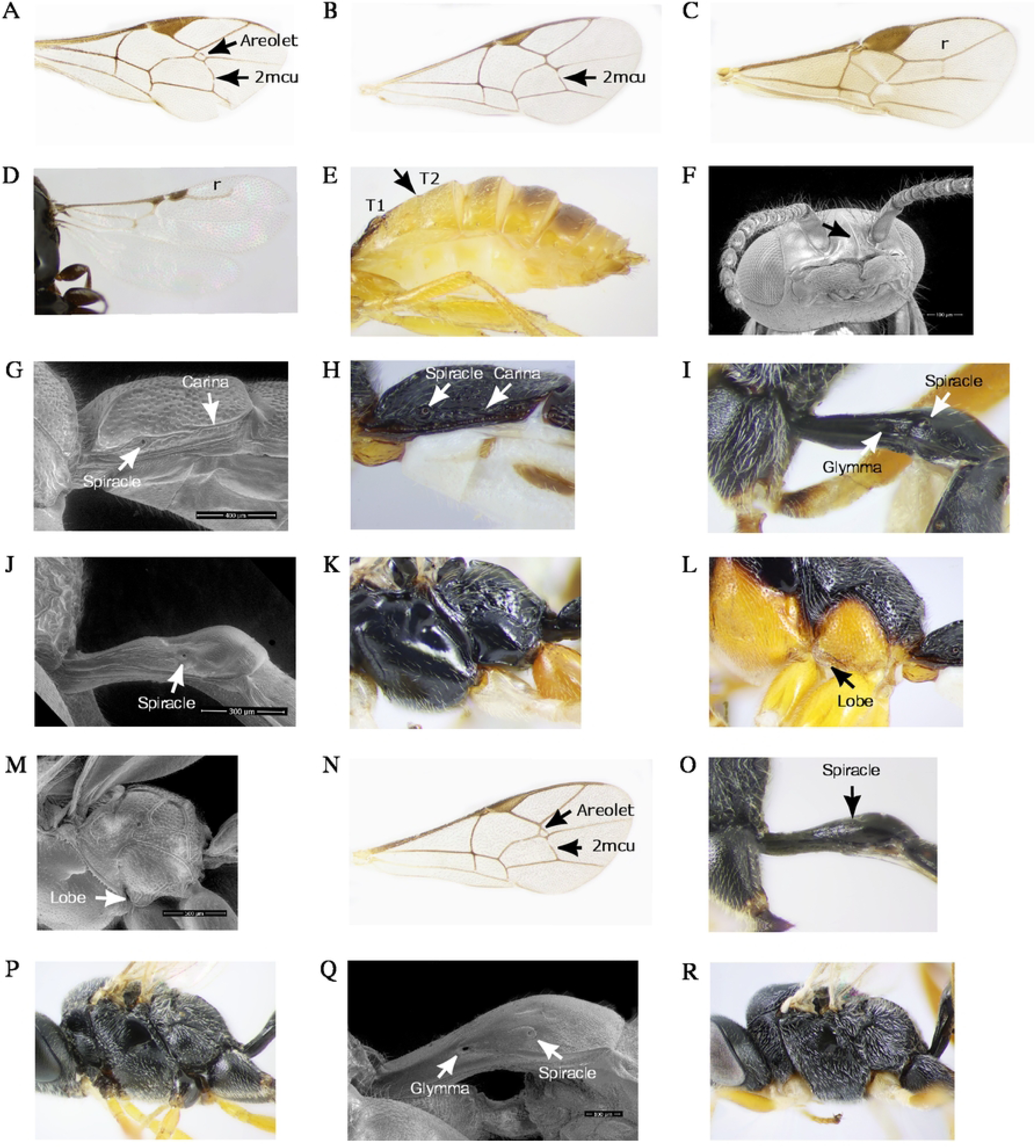
Key morphological characters for the identification of the GBM larval parasitoids found in northwest Pennsylvania. **(A)** subtriangular areolet and vein 2m-cu, **(B)** absent areolet with vein 2m-cu present, **(C)** enclosed radial cell, **(D)** open radial cell, **(E)** 2^nd^ and 3^nd^ tergites fused, **(F)** clypeus with a strong angular or subangular median lobe, **(G-H)** spiracle of 1^st^ tergite with a complete dorsolateral carina, **(I and Q)** spiracle of the first abdominal tergite placed posteriorly and glymma, **(J and O)** spiracle of first abdominal tergite with absent glymma, **(K)** carina forming a lobe, **(L-M)** submetapleura carina extending anteriorly forming a lobe, **(N)** closed areolet before the junction with radius, **(P)** black coxae, **(R)** yellow fore and mid coxae.

1b. Fore wing vein 2m-cu absent; hind wing venation reduced (Figs 5C-D). 2

2a. Face and clypeus separated by a groove, mandibulates sometimes exodont; 2^nd^ and 3^nd^ tergites usually fused (Fig 5E); predominantly yellow coloration; radial cell completely enclosed (Fig 5C)… Braconidae (*Bracon variabilis;* Figs 3A-B).

2b. Not agreeing with the above; clypeus with a strongly produced angular or subangular median lobe (Fig 5F); predominantly black coloration; radial cell not enclosed (Fig 5D). Bethylidae (*Goniozus fratellus;* Figs 3C-D).

3a. Spiracle of 1^st^ tergite placed behind or around the middle of the tergite (Figs 5G-H) ……………………………………………………………………………………………………..4 3b. Spiracle of 1^st^ tergite placed at posterior third of the tergite (Figs 5I and Q) …….……..……5

4a. Areolet present and subtriangular, wider than high (Fig 5A); nervellus intercepted below the middle or in some cases in the middle; submetapleura carina not extending anteriorly forming a lobe (Fig 5K). Ichneumonidae (*Scambus*; Figs 3E-F).

4b. Areolet absent (Fig 5B); nervellus intercepted near hind end the middle; dorsolateral carina of the first tergite complete (Fig 5G); strong submetapleura carina extending anteriorly forming a lobe (Figs 5L-M). Ichneumonidae (*Glypta*; Figs 3G-H).

5a. Areolet present and closed before the junction with radius (Fig 5N); glymma absent (Figs 5J and O); all coxae black (Fig 5P). Ichneumonidae (*Campoplex tortricidis*; Figs 3I-J).

5b. Areolet absent (Fig 5B); glymma present (Figs 5I-Q); fore and mid coxae yellow (Fig 5R)… Ichneumonidae (*Enytus obliteratus*.; Figs 3K-L).

## Discussion

The GBM, *P. viteana* has an abundant number of larval parasitoids that provide significant natural control. We successfully identified eight hymenopteran parasitoids in samples from six commercial Concord vineyards in northwest Pennsylvania over two consecutive years (2023 and 2024). These vineyards were conventionally managed following current spray recommendations [30], indicating that natural control still occurs despite insecticide use. Field parasitism rates varied through the growing season and ranged from 0 - 39% in 2023 and from 0 - 52.1% in 2024. This project represents a first step toward our understanding of the native natural enemies present in the Lake Erie Region and their use in pest management programs.

The community of larval parasitoids identified in this study confirms previous species records and provides new findings. *E. obliteratus*, *C. tortricidis, B. variabilis,* and *Scambus* sp. have been previously reported parasitizing *P. viteana* [4,5,10], but *G. fratellus, G. depressa* cf, *G. ohioensis* cf, and *G. ignota* are new reports for this species. In our samplings, the most abundant parasitoid species was *B. variabilis* accounting for 56.6 % (in 2023) and 68.6% (in 2024) of all parasitoids found. However, in similar studies conducted in 1986-87 in the Finger Lakes in New York, the most abundant species was the egg parasitoid *T. pretiosum*, followed by the larval parasitoids *A. polychrosidis,* and *G. mutica* from the eight species found [4]. In a 3-year (2003- 2005) study in Michigan vineyards, *Sinophorus* sp. was the most abundant parasitoid, comprising 11-76% of all the specimens collected from 12 identified species [5]. These results confirm the presence of a large number of GBM parasitoids in the eastern U.S. and suggest that the community composition and abundance of these species vary in different grape-growing regions.

The field parasitism rates fluctuated over the growing season and habitat. In 2023, the highest percent of parasitism was from samples collected in wild grapes early in the season and ranged from 0 - 71.4% (Fig 2A; S1 Table). Similar results were found by Seaman et al. [4], who reported higher parasitism in wild grapes reaching up to 65%. The highest parasitism rates in cultivated grapes were found during the first week of August and ranged from 4.5 - 22.2% in 2023 and from 2 - 52.1% in 2024 (Fig 2; S1 and S2 Tables). In contrast, the lowest percentages of parasitism were found during the second week of July 2023 and the last week of June 2024. Similarly, Jenkins et al. [31] found a low abundance of parasitoids early in the season that increased in mid-August to levels of up to 30% and declined as the season progressed. In the Lake Erie region, the first generation of GBM emerges before the cultivated grapes bloom and feeds on developing blossoms from wild grapes. In comparison to later generations that burrow into grape berries, these larvae are highly exposed to parasitoids and predators despite building protective silk enclosures. Gleisner [10] suggested that berries may protect the larva from parasites with short ovipositors. The variation in parasitism through the season could be associated with varying availability of appropriate GBM instars to use for parasitoids, variation in GBM infestation levels, or asynchrony between parasites and the host [5,10]. Other possibilities include changes in the availability of alternate hosts for parasitoids or differences in the timing of diapause initiation for the parasitoids and the host later in the season.

All the GBM parasitoids found in this study have alternate hosts. The most abundant parasitoid species in our samplings was the ectoparasitoid *B. variabilis,* whose parasitism rates ranged from 1.3 - 37.3% in 2023 and from 1.6 - 51.1% in 2024. This species was previously reported parasitizing GBM in Van Buren and Berrien Counties in Michigan and in the Finger Lakes region in NY [4,5]. Other lepidopteran hosts of *B. variabilis*, include the nantucket pine tip moth, *Rhyacionia frustrana* (Lepidoptera: Tortricidae) in Georgia [32], the hickory shuckworm moth, *Cydia caryana* (Lepidoptera: Tortricidae) in Texas [33], and the alder tubemaker moth, *Acrobasis rubrifasciella* (Lepidoptera: Pyralidae) in Eaglenest Lakes, near Ely, Minnesota [34].

*B. variabilis* has also been reported parasitizing the curculionid flower beetle, *Baris transversa* [35]. The endoparasitoid *E. obliteratus* was the second most abundant species with parasitism rates in commercial vineyards ranging from 1.3 - 9.4% in 2023 and from 1.5 - 16.3% in 2024. This species has been previously reported parasitizing GBM in Michigan State [5]. *E. obliteratus* also parasitizes larvae of the European GBM, *Lobesia botrana* [36], and larva of the oriental fruit moth, *Grapholita molesta*, an important pest of stone fruits (peaches and apples, cherries). It has also been reported as a parasitoid of *Arogalea cristifasciella* (Lepidoptera: Gelechiidae) larva [37]. This genus has primarily a Nearctic and Palearctic distribution [38]. The ectoparasitoid *G. fratellus* had parasitism rates that fluctuated from 5.4 - 7.7% in 2023 and from 1 - 11.2% in 2024. To our knowledge, this is the first time this species has been reported preying on GBM. *G. fratellus* also parasitizes *Acrobasis nuxvorella* (Neunzig) (Lepidoptera: Pyralidae) [33]. This species has been reported in several States, including California, Nevada, Arizona, New Mexico, Georgia, Texas, Florida, South Carolina, Michigan, Connecticut, Massachusetts, Ontario, and Louisiana [23] without information about the hosts. The endoparasitoid ***C. tortricidis*** had parasitism rates below 2%. This species was first found by Cushman [20] in North East, PA parasitizing *P. viteana,* and was reported again by Gleissner [10] from GBM samples gathered between 1940-1943 in the same geographic location. It was also reported parasitizing the oriental fruit moth in Ohio [39]. Information on this species is scarce and the only other geographical record besides northwest PA is the one in Edmonton, Canada [40,41].

There were also three species of *Glypta* identified in this study: *G. depressa*, *G. ohioensis*, and *G. ignota which* together had field parasitism rates below 5%. To our knowledge, this is the first report of these species parasitizing GBM and no other hosts have been reported in the literature.

Information on their geographic distribution is also limited. *G. depressa* has been recorded in New Mexico, whereas *G. ohioensis* has been found in Ohio, Louisiana, Michigan, New York, Pennsylvania, South Carolina, West Virginia and Ontario [21]. For *G. ignota*, there is only one report in Nova Scotia [21]. Other species of this genus, such as *G. animosa*, *G. vulgaris*, and *G. mutica* are known to parasite *P. viteana* [5,9]. Natural parasitism rates of *G. mutica* in the Finger Lakes ranged from 0.01 - 6.4% and in caged experiments varied from 3-7% [4]. Glypta species in general, are known parasites of several Lepidoptera families, including Cochylidae, Gelechiidae, Geometridae, Lasiocampidae, Lycaenidae, Lymantriidae, Momphidae, Noctuidae, Pyralidae and Tortricidae. This genus is found principally in the Holarctic and is less frequent in the Neotropic and Oriental regions.

*Scambus sp.* was a less frequent parasitoid found in our study only in the 2023 samplings with parasitisms lower than 1%. Two species within this genus have been previously identified parasitizing *P. viteana*a: *S. brevicornis* and *S. hispae* in Michigan vineyards [5]. However, other *Scambus* species have been reported parasitizing other tortricids. For example, *Scambus elegans* parasitize *L. botrana* [42], whereas *Scambus calobatus* (Gravenhorst) and *Scambus vesicarius* (Ratzeburg) parasitize the European grapevine moth, *Eupoecilia ambiguella* (Lepidoptera: Tortricidae) [43–45]. There are also reports of *Scambus* species parasitizing Diptera and Coleoptera insects [46–48]. This genus is widely distributed in the Northern Hemisphere [19].

The abundance of the GBM parasitoids found in this study fluctuated over the course of the growing season. *E. obliteratus, C. tortricidis* and *Glypta sp.* were only present from June to mid-August, whereas *B. variabilis* and *G. fratellus* were found from late July to September (Fig 2).

This temporal variation could be associated with different thermal requirements and diapause periods for the different species and could be helpful in reducing interspecific competition of parasitoids for the same prey.

We conclude that the American GBM, *P. viteana* is host for several larval parasitoids in commercial Concord vineyards in Pennsylvania. We confirmed previous species reports and added four new records to the parasitoid list of this species. The parasitoid abundance differed over the season but was greatest in early August reaching parasitism rates of up to 52.1%.

Despite their importance, information about these parasitoids is limited; future research is needed to document the biology and ecology of the most promising species to identify ways to increase their natural populations in vineyards. Measurements should be taken to protect the community of natural enemies that prey on GBM in vineyards. Some strategies, include reducing the use of broad-spectrum insecticides, using economic thresholds to determine the need for insecticide applications, and protecting wild vegetation that provides food for parasitoid adults.

## Acknowledgments

We would like to thank the six farmers who graciously allowed us to conduct our parasitoid samplings at their vineyards. Many thanks to Laura Porturas, Assistant curator of the Frost Entomological Museum; Jacki Whisenant, Insect Technician of the Cornell University Insect Collection; and Dr. Robert Kula, coordinator of the USDA Systematic Entomology Laboratory for graciously allowing us to visit and inspect the entomological collections to confirm the parasitoid identifications. We are very thankful to Mr. Jerry Magraw for taking the SEM images of some parasitoid specimens and structures and to Miss Norah Dana for logistic support. Thanks to the Lake Erie Regional Grape Research and Extension Center for providing grape clusters for insect rearing. Lastly, special thanks to the anonymous reviewers for their time and valuable suggestions.

grant number: 162873

## Author contributions

Conceptualization, Methodology, Visualization, and Validation: Flor E. Acevedo and Jesus H. Gomez-Llano

Data curation: Neetu Khanal, Jesus H. Gomez-Llano and Flor E. Acevedo

Formal Analysis, Funding Acquisition, Project Administration, supervision: Flor E. Acevedo

Investigation: Neetu Khanal, Jesus H. Gomez-Llano and Flor E. Acevedo

Writing – Original Draft Preparation, review, and editing: Flor E. Acevedo, Jesus H. Gomez-Llano, and Neetu Khanal

## Data Availability Statement

All relevant data are within the manuscript.

## Supporting information

**S1 Table.** GBM larval parasitism in field conditions per sampling site throughout the 2023 growing season.

**S2 Table.** GBM larval parasitism in field conditions per sampling site throughout the 2024 growing season.

